# Pynoma, PyABraOM and BIOVARS: Towards genetic variant data acquisition and integration

**DOI:** 10.1101/2022.06.07.495190

**Authors:** Paola Carneiro, Felipe Colombelli, Mariana Recamonde-Mendoza, Ursula Matte

## Abstract

**Motivation:** Advances in genomic sequencing of human populations have generated a large amount of genomics data deposited in multiple sources. Programmatic batch searches executed at once are of great scientific interest to ease genomic investigations by retrieving and integrating this massive and decentralized data with little manual intervention.

**Results:** Pynoma and PyABraOM APIs were developed to offer multiple queries in gnomAD and ABraOM databases, respectively. A centralized search in these databases with data integration is offered by a third API, BIOVARS, which combines the resulting information with statistical and graphical visualizations. The implemented features are demonstrated in a case study using *ACE2, ADAM17* and *TMPRSS2* genes, which presents a generalizable workflow that shows how our APIs facilitate the access and integration of valuable biological data.

**Availability:** All the APIs are written in Python 3. Graphical visualizations for the retrieved data are provided by using the R language version 4.1. The source codes are publicly available and hosted on GitHub (github.com/bioinfo-hcpa).

**Supplementary information:** Supplementary data is available at github@nbioinfo-hcpa.

## 1 Introduction

Modern sequencing technologies have generated a vast amount of genome data and have transformed biology into a data-rich discipline, bringing numerous scientific benefits. In human genetics, these advances aid in molecular diagnostics and ancestry studies by comprehensively mapping the diverse human genetic variations across populations. However, the circumstances of the growing genetic information demand an easy and programmatic approach to obtain data for multiple searches in one or multiple different databases.

The development of distinct genetic variant databases helps store and share data on genetic variability across the world. Earlier bona fide efforts of gnomAD (Karczewski *et al*., 2020) and ABraOM (Naslavsky *et al*., 2022) databases are excellent examples of how these resources assist the scientific community and clinicians in studying human genomics diversity and identifying disease-causing variants. Currently, AbraOM and gnomAD include genetic variation information among 1171 and 76156 genomes from Whole Genome Sequence for GRCh38/hg38. To note, ABraOM provides a source of Brazilian genetic variation that is underrepresented in gnomAD and 2 million previously unknown variants in public databases (Naslavsky *et al*., 2022). However, these databases are not suitable for performing multiple queries or querying multiple databases simultaneously. This limitation may cause the delay of research projects since manually retrieving all the needed information can be an exhaustive and time consuming process. To overcome this challenge, software solutions like application programming interfaces (APIs) may be created to help researchers handle information for multiple queries. A particularly interesting feature of these APIs emerges when just one configuration of inputs and little programming experience is necessary to retrieve the target information from different databases.

Here, we propose two APIs, Pynoma and PyABraOM, that provide access and return detailed information on population genetic variants from gnomAD and ABraOM databases, respectively. Additionally, to address the gap of easily combining information from both resources in population genetic studies, we develop a third API called BIOVARS that provides a centralized programmatic approach to query and merge genetic variant information from gnomAD and ABraOM.

## 2 Implementation

The APIs’ core functionalities were built and tested on Python 3. Scripts implemented in R v4.1 provide graphical visualizations for the retrieved data, which are invoked and interfaced by BIOVARS’s Plotter class. Among the proposed visualizations, there are static plots and an interactive variant report generated using HTML and CSS.

Four kinds of searches are supported by the APIs to return data from gnomAD and ABraOM databases. Both APIs accept searches by gene, genome localization, and specific variants. However, due to differences in databases’ scope, searches by transcript and the gnomAD’s variant IDs are only available for the Pynoma API. The server requests can be performed using the two latest human genome versions, GhRC37/hg19 and GhRC38/hg38.

To integrate the data coming from the two databases, the gnomAD’s variant ID attribute was used. This ID is a string composed of the chromosome and the genomic region where the variant is located, as well as the reference and alternative alleles. This way, it is possible to create an unique ID that acts as an indexer for the integration. For data derived from ABraOM database, the variant ID is built using the information from the resulting request when the variant is not an indel. However, for indel variants, additional information has to be retrieved from a source such as the Ensembl database. This process can consume significantly more time, but it is necessary for indexing the variants and properly integrate the data.

To allow easy further integration with different databases, BIOVARS was built using the Object-Oriented programming paradigm. There is a specific class, *Sources*, responsible for validating the types of searches depending on which databases are selected by the user for searching the variants from. If a new database is integrated with its particular types of searches into BIOVARS, it is possible to validate the search call through this class without hindering the code’s legibility. Additionally, *Sources* provides a way to validate other input parameters that may require special attention, such as the accepted genome versions of each database.

However, it is important to clarify that all the requests to the databases are implemented by specific APIs designed to deal with the particularities of these servers. BIOVARS only acts as a bridge, providing useful methods to integrate the different information and data formats. Additionally, these database-specific APIs are self-contained and have additional features, ideal to be used alone if the user only wants to work with a particular database. This is why our proposal is divided into three different APIs: one for retrieving data from ABraOM (PyABraOM), the other for retrieving data from gnomAD (Pynoma), and the last one for integrating the multi-source variant data (BIOVARS).

## 3 Discussion

The standard workflow for acquiring the desired data about genetic variants requires the user to navigate to each specific database website and interact with the search boxes and other user interface components. If a particular user desires to collect variants information of various genes, then a manual query is necessary for each gene, followed by data processing to save results. Not only this manual process could be very time-consuming and error-prone, but it would also make bioinformatics pipelines impossible to be fully automated if a procedure inside them generates the input used in a genetic variants search that is required for completing the pipeline’s execution. The described problem is fully solved by using our proposed APIs, which demand little programming knowledge.

Fig. 1A demonstrates how the developed APIs allow easy and effortless search for variants in an arbitrary list of genes using gnomAD, ABraOM or both databases simultaneously. A partial view of the table returned by the BIOVARS integrated search is provided in Fig. 1B. The results tables generated by our APIs are given in pandas DataFrame objects and can be easily exported if necessary (see pandas documentation^1^).

**Fig. 1:**
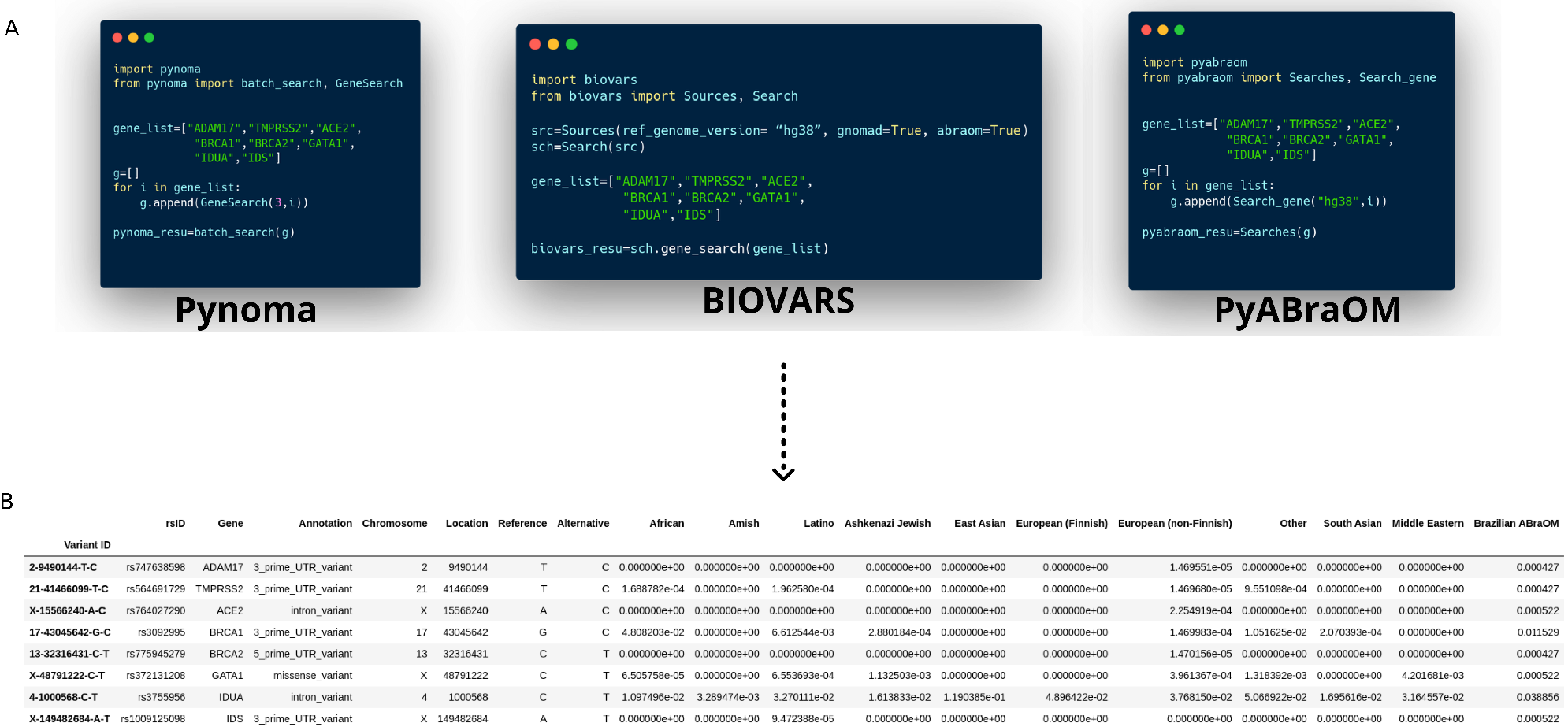
(A) Query demonstration for a list of selected genes for each API and (B) an example of results table returned by BIOVARS query containing merged information of gnomAD and ABraOM.

Besides the data automation possibility and the multi-source data integration, our proposed tools also allow plotting the results using different visualization schemes provided by BIOVARS. Fig. 2 illustrates some of the implemented visualizations for the variants found in both databases for the *ACE2* gene. To use these visualizations with query results from Pynoma or PyABraOM retrieved data, the resulting tables must be converted to the same format used by BIOVARS with the appropriate methods implemented by the BIOVARS’ Search class. The case study described in the supplementary material discusses in more detail the workflow for retrieving data with the APIs and how to explore the visualization options provided.

**Fig. 2:**
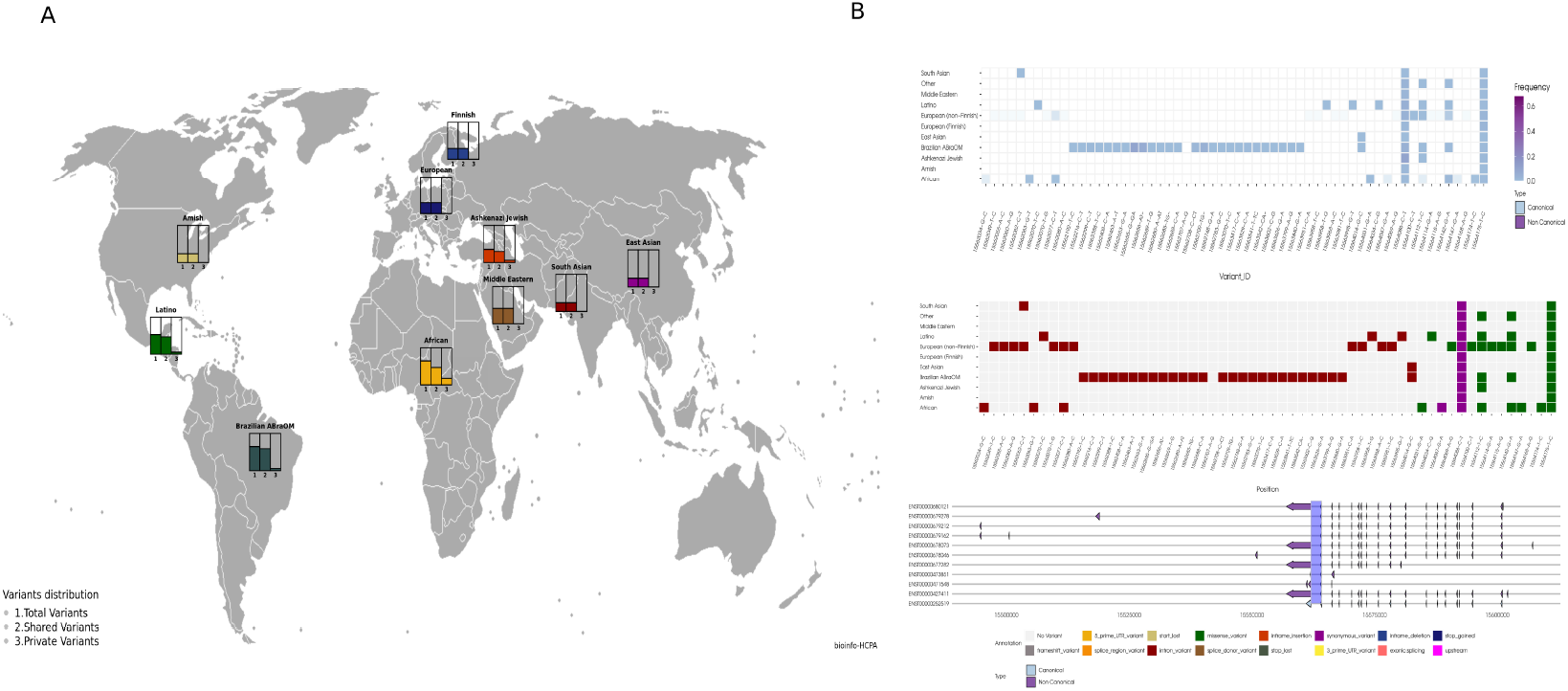
Query results for gene *ACE2* using BIOVARS. (A) Number of total, shared, and private variants among different human populations. (B) The genomic localization of *ACE2* variants is provided according to the allele frequency of each population (upper graph), variant annotation (middle graph) and transcript orientation (bottom graph).

## 4 Case Study

Variants in the genes *ACE2, TMPRSS2*, and *ADAM17* have been reported to modulate SARS virus infections (Hou *et al*., 2020; Lambert *et al*., 2005). For example, rare genetic variants were identified in patients with severe cases of COVID-19 in a small Russian cohort, and were likely to predispose to severe COVID-19 infection due to upshift inflammatory response in these patients (Shikov *et al*., 2020). In this setting, human genetic databases are valuable resources to recognize the genetic variants and their frequencies across human populations for estimating disease susceptibility.

To illustrate the functionalities of the proposed tools, a case study using the genes *ACE2, TMPRSS2* and *ADAM17* was performed. Results for all three genes comprised over 2,000 variants and BIOVARS output shown in figure 1 for *ACE2* (similar to the structure shown in Fig. 1) was obtained in less than one hour. Detailed results are provided in the supplementary file. With this case study, our goal is to demonstrate a generalizable workflow that reinforces the easy access and integration of valuable genomic data using our APIs.

## 5 Conclusion

Tackling the challenge of easily accessing large amounts of human genomic variants data from databases such as gnomAD and ABraOM is of particular interest to large-scale genomic studies and clinical practice.

Additionally, for integrating the acquisition and processing of such data into bioinformatics pipelines and tools, it is imperative to automate this access, reducing the need for human intervention. This paper proposes Pynoma and PyABraOM APIs to easily automate data acquisition from gnomAD and ABraOM databases. We also provide a third API, BIOVARS, with which users can readily integrate the data retrieved from these two different databases and plot them in different informative visualizations.

## Footnotes

1 https://pandas.pydata.org/docs/

## References

Hou, Y., Zhao, J., Martin, W., Kallianpur, A., Chung, M. K., Jehi, L., Sharifi, N., Erzurum, S., Eng, C., and Cheng, F. (2020). New insights into genetic susceptibility of covid-19: an ace2 and tmprss2 polymorphism analysis. BMC medicine, 18(1), 1–8.

Karczewski, K. J., Francioli, L. C., Tiao, G., Cummings, B. B., Alföldi, J., Wang, Q., Collins, R. L., Laricchia, K. M., Ganna, A., Birnbaum, D. P., et al. (2020). The mutational constraint spectrum quantified from variation in 141,456 humans. Nature, 581(7809), 434–443.

Lambert, D. W., Yarski, M., Warner, F. J., Thornhill, P., Parkin, E. T., Smith, A. I., Hooper, N. M., and Turner, A. J. (2005). Tumor necrosis factor-α convertase (adam17) mediates regulated ectodomain shedding of the severe-acute respiratory syndrome-coronavirus (sars-cov) receptor, angiotensin-converting enzyme-2 (ace2). Journal of Biological Chemistry, 280(34), 30113–30119.

Naslavsky, M. S., Scliar, M. O., Yamamoto, G. L., Wang, J. Y. T., Zverinova, S., Karp, T., Nunes, K., Ceroni, J. R. M., de Carvalho, D. L., da Silva Simões, C. E., et al. (2022). Whole-genome sequencing of 1,171 elderly admixed individuals from são paulo, brazil. Nature communications, 13(1), 1–11.

Shikov, A. E., Barbitoff, Y. A., Glotov, A. S., Danilova, M. M., Tonyan, Z. N., Nasykhova, Y. A., Mikhailova, A. A., Bespalova, O. N., Kalinin, R. S., Mirzorustamova, A. M., et al. (2020). Analysis of the spectrum of ace2 variation suggests a possible influence of rare and common variants on susceptibility to covid-19 and severity of outcome. Frontiers in genetics, 11.

