## Supplementary file for "Pynoma, PyABraOM and BIOVARS: Towards genetic variant data acquisition and integration"

BIOINFORMATICS CORE
HOSPITAL DE CLÍNICAS DE PORTO ALEGRE

### Pynoma, PyABraOM and BIOVARS: Towards human genetic variant data acquisition and integration

*Paola* CARNEIRO[a,b]*, *Felipe* COLOMBELLI[c]*,
*Mariana* RECAMONDE-MENDOZA[a,c] *and Ursula* MATTE[a,b,d,+]

[a]*Bioinformatics Core, Hospital de Clínicas de Porto Alegre , Porto Alegre, 990035-903, Brazil.*
[b]*Graduate Program in Genetics and Molecular Biology, Universidade Federal do Rio Grande do Sul, 91501-970, Brazil.*
[c]*Institute of Informatics, Universidade Federal do Rio Grande do Sul, Porto Alegre, 91501-970, Brazil.*
[d]*Departament of Genetics, Universidade Federal do Rio Grande do Sul, 91501-970, Brazil.*
[*]*These authors contributed equally to this work.*
[+]*Reference Contact:*

June 22, 2022

**References** **20**

CHAPTER 1

### About

This documentation in the format of case study refers to the unpublished version of BIOVARS, Pynoma and PyABraOM APIs. Thus, the reader must be aware of possible future modifications, which will be tracked in the projects' repositories.

The presented APIs are Python v3 packages, containing R v4 scripts (BIOVARS). They are freely available on the Bioinformatics Core of Hospital de Clínicas de Porto Alegre (NBioinfo-HCPA) GitHub organization profile (https://github.com/bioinfo-hcpa), and currently only published on GitHub under the GPL-3.0 License (refer to the repositories). All data tables and the HTML file generated in this case study are publicly available on GitHub.

The presented packages intend to provide an easy way to access and integrate data from gnomAD and ABraOM databases. They are under active development by researchers at NBioinfo-HCPA and will provide support for integrating and accessing more human genetic variant databases in the future. As open source projects, we invite anyone interested in the tools to contribute for their improvement and upgrades.

### CHAPTER 2

#### Installation

All the packages are publicly available and hosted on the Github repository of the Nbioinfo-HCPA[1]. Some libraries are required to properly use these packages, however all these libraries are already defined as requirements for the installation of the APIs through the *requirements.txt* file, so no further installation is needed for their setup regarding Python, although, for using the R functions, additional configuration is necessary.

Note that, despite Pynoma and PyABraOM being independent on each other, BIOVARS requires both packages to be installed. For installing the latest version of the packages, download them from their GitHub repositories. For installing the latest stable version, simply install them from PyPI with a *pip* command.

```
# Install latest version of the packages
$ git clone https://github.com/bioinfo-hcpa/pynoma.git
$ git clone https://github.com/bioinfo-hcpa/pyabraom.git
$ git clone https://github.com/bioinfo-hcpa/biovars.git
$ pip install -e pynoma
$ pip install -e pyabraom
$ pip install -e biovars

# Install the latest stable version
$ pip install pynoma
$ pip install pyabraom
$ pip install biovars
```

BIOVARS also provides different graphical visualization methods for the retrieved data. All the visualizations were made using R tools, and the R scripts for generating these visualizations are called from the Python code using *rpy2* Python-R bridge library. However, the R required packages are not automatically set up with the pip command and, thus, should be manually installed through R shell, as follows.

[1]https://github.com/bioinfo-hcpa/

```
$ R
> pkgs <- c("ggplot2", "ggthemes", "gridExtra", "egg", "png", "grid",
    "cowboy", "patchwork", "httr", "jsonlite", "xml2", "dplyr",
    "RColorBrewer", "stringr", "gggenes")
> install.packages(pkgs)
```

### CHAPTER 3

### Case Study

The *ACE2* gene (OMIM #300335) encodes angiotensin converting enzyme 2 receptor that allows the attachment SARS-CoV-2 with spike glycoprotein (S-protein) and promotes fusion of virus in host cells. On the other hand, *TMPRSS2* (OMIM #602060) encodes a transmembrane protease serine 2 which promotes the virus entry by the proteolytic cleavage of *ACE2* after binding to the S-protein. *ADAM17* (OMIM #603639) is an ectodomain "sheddase" that performs modulation of *ACE2* protein and immune surveillance in response to pathogen infections. The interplay of these proteins were addressed in the literature related to the viral S-protein with *ACE2*-*TMPRSS2* (Glowacka et al., 2011) and the regulation of *ACE2* protein shedding by *ADAM17* (Lambert et al., 2005).

APIs can provide an interface for connecting with human genetic databases and retrieve data to further explore variants in these genes. Such programs can then be extended to build a complete pipeline with additional functionalities enabled by the accessed data. In our implemented APIs, this access in each database is handled by Python packages created to accept different types of inputs for searching data in one or more databases at once. Here, we will demonstrate how to retrieve data using multiple batch search functions provided by our packages to retrieve data from gnomAD (Karczewski et al., 2020) and ABraOM (Naslavsky et al., 2022) databases. To demonstrate a workflow of the package functionalities, Pynoma and PyABraOM use only *ADAM17* and *TMPRSS2* genes, while BIOVARS uses all of the selected genes in this case study.

#### 3.1 Pynoma

Once the libraries are installed, the first step is to import the Pynoma modules for the desired query types. Optional parameters can be set by the user when calling the *get_data()* or *batch_search()* methods. For example, *additional_population_info* that includes information about the allele frequency for each population in gnomAD (see the documentation in the repository for more information[1]).

[1] https://github.com/bioinfo-hcpa/pynoma

The gene query provides data that comprehends the variants' annotation, sequencing methodologies, allele frequencies and allele number. In the code example below, the *pynoma_df_1* object is a table with 25 columns, containing information for 2123 variants according to the hg38 genome version using the Pynoma batch search functionality. Some details of this table are shown in Figure 3.1.

```
# Load package and functions for gene, region, variant and batch search
from pynoma import GeneSearch, RegionSearch, TranscriptSearch,
    VariantSearch, batch_search

# Prepare inputs of  different search types
gene_list = [GeneSearch(3,"ADAM17"), GeneSearch(3,"TMPRSS2")]

region_list = [RegionSearch(3, (2,9490303,9493719)),
               RegionSearch(3, (21,41480500,41480570))]

t_list = [TranscriptSearch(3, "ENST00000310823"),
          TranscriptSearch(3, "ENST00000332149")]

mixed_list = gene_list + region_list + t_list

# Batch search
# Search in gnomAD database by gene
pynoma_df_1 = batch_search(gene_list, additional_population_info=True)

# Search in gnomAD database by regions
pynoma_df_2 = batch_search(region_list, additional_population_info=True)

# Search in gnomAD by transcript ID
pynoma_df_3 = batch_search(t_list, additional_population_info=False)

# Search in gnomAD for different queries
pynoma_df_4 = batch_search(mixed_list, additional_population_info=True)
```

| | Variant ID | Gene | Annotation | Allele Count | Allele Number | Allele Frequency | Number of Homozygotes | Chromosome | Location | Reference |
|---|---|---|---|---|---|---|---|---|---|---|
| **0** | 2-9490106-C-T | ADAM17 | 3_prime_UTR_variant | 1 | 108646 | 0.000009 | 0 | 2 | 9490106 | C |
| **1** | 2-9490111-ATTGAT-A | ADAM17 | 3_prime_UTR_variant | 560 | 108994 | 0.005138 | 2 | 2 | 9490111 | ATTGAT |
| **2** | 2-9490112-T-C | ADAM17 | 3_prime_UTR_variant | 9 | 107922 | 0.000083 | 0 | 2 | 9490112 | T |
| **3** | 2-9490115-AT-A | ADAM17 | 3_prime_UTR_variant | 77991 | 151428 | 0.515037 | 20911 | 2 | 9490115 | AT |
| **4** | 2-9490117-TTGTA-T | ADAM17 | 3_prime_UTR_variant | 1 | 37368 | 0.000027 | 0 | 2 | 9490117 | TTGTA |
| **2118** | 21-41508071-C-A | TMPRSS2 | 5_prime_UTR_variant | 1 | 152036 | 0.000007 | 0 | 21 | 41508071 | C |
| **2119** | 21-41508074-C-G | TMPRSS2 | splice_region_variant | 1 | 152150 | 0.000007 | 0 | 21 | 41508074 | C |
| **2120** | 21-41508074-C-T | TMPRSS2 | splice_region_variant | 1 | 152150 | 0.000007 | 0 | 21 | 41508074 | C |
| **2121** | 21-41508075-G-C | TMPRSS2 | splice_region_variant | 4 | 151968 | 0.000026 | 0 | 21 | 41508075 | G |
| **2122** | 21-41508076-C-G | TMPRSS2 | splice_region_variant | 1 | 152170 | 0.000007 | 0 | 21 | 41508076 | C |

| | Alternative | African | Amish | Latino | Ashkenazi Jewish | East Asian | European (Finnish) | European (non-Finnish) | South Asian | Middle Eastern |
|---|---|---|---|---|---|---|---|---|---|---|
| **0** | T | 0.000000e+00 | 0.000000e+00 | 0.000000e+00 | 0.000000e+00 | 0.000000e+00 | 0.000000e+00 | 2.274899e-05 | 0.000000e+00 | 0.000000e+00 |
| **1** | A | 2.687656e-03 | 0.000000e+00 | 8.415033e-03 | 2.846300e-03 | 0.000000e+00 | 1.225190e-04 | 7.901747e-03 | 1.321353e-03 | 5.952381e-03 |
| **2** | C | 6.432937e-05 | 0.000000e+00 | 0.000000e+00 | 0.000000e+00 | 1.944012e-04 | 0.000000e+00 | 9.170526e-05 | 5.285412e-04 | 0.000000e+00 |
| **3** | A | 4.816798e-01 | 7.285714e-01 | 4.157637e-01 | 6.279607e-01 | 9.118892e-02 | 4.654291e-01 | 5.928006e-01 | 4.359718e-01 | 6.687898e-01 |
| **4** | T | 8.981498e-05 | 0.000000e+00 | 0.000000e+00 | 0.000000e+00 | 0.000000e+00 | 0.000000e+00 | 0.000000e+00 | 0.000000e+00 | 0.000000e+00 |
| **2118** | A | 2.414409e-05 | 0.000000e+00 | 0.000000e+00 | 0.000000e+00 | 0.000000e+00 | 0.000000e+00 | 0.000000e+00 | 0.000000e+00 | 0.000000e+00 |
| **2119** | G | 0.000000e+00 | 0.000000e+00 | 0.000000e+00 | 0.000000e+00 | 0.000000e+00 | 0.000000e+00 | 1.470675e-05 | 0.000000e+00 | 0.000000e+00 |
| **2120** | T | 0.000000e+00 | 0.000000e+00 | 0.000000e+00 | 0.000000e+00 | 0.000000e+00 | 0.000000e+00 | 1.470675e-05 | 0.000000e+00 | 0.000000e+00 |
| **2121** | C | 0.000000e+00 | 0.000000e+00 | 0.000000e+00 | 0.000000e+00 | 0.000000e+00 | 0.000000e+00 | 5.886162e-05 | 0.000000e+00 | 0.000000e+00 |
| **2122** | G | 0.000000e+00 | 0.000000e+00 | 0.000000e+00 | 0.000000e+00 | 1.928268e-04 | 0.000000e+00 | 0.000000e+00 | 0.000000e+00 | 0.000000e+00 |

Figure 3.1: Pynoma gene query batch search result for *ADAM17* and *TMPRSS2* on gnomAD database. For better visualization, a partial view of results contained the first and last five rows of 19 columns out of 25 from the data table for *ADAM17* and *TMPRSS2*, respectively, is demonstrated.

#### 3.2 PyABraOM

After the installation (see Chapter 2), import the functions from pyABraOM package for the desired query types. Additional parameters such as variant identifier (rsID) and CEGH filter can be set. Another possibility is to select only variants that passed in GATK filter (see the documentation in the repository for more information[2]).

The gene search provides data that comprehends the variants' annotation, variant quality filters, allele frequencies and allele number. In the code example below, the *pyabraom_df_1* object is a table with 13 columns, containing information for 2744 variants according to the hg38 genome version using the PyABraOM batch search functionality. Some details of this table are shown in Figure 3.2.

```
# Load package and functions for Gene, Region, Variant and Batch search
from pyabraom import Search_gene,Search_region,Variant_ID,Searches

# Prepare inputs of  different search types
gene_list = ["ADAM17","TMPRSS2"]
region_list = [(2,9490303,9493719) ,(21,41480500,41480570)]
rsIDs=["rs140298748","rs576215861"]

# Batch search
# Search on ABraOM database by gene
pyabraom_df_1 = Searches([Search_gene("hg38",gene_list[0],
                          Variant_ID=True,CEGH_Filter=True),
                         Search_gene("hg38",gene_list[1],
                         Variant_ID=True,CEGH_Filter=True)])

# Search on ABraOM database by regions
pyabraom_df_2 = Searches([Search_region("hg38",region_list[0]),
                         Search_region("hg38",region_list[1])])

# Search on ABraOM database by rsID
pyabraom_df_3 = Searches([Variant_ID("hg38",rsIDs[0]),
                         Variant_ID("hg38",rsIDs[1])])
```

| | Chromosome | Position | Reference | Alternative | Annotation | Gene | rsID | GATK Filter | CEGH Filter | Number of Homozygotes | Allele Number | Allele Count | Allele Frequency |
|---|---|---|---|---|---|---|---|---|---|---|---|---|---|
| **0** | 2 | 9488500 | TATT | - | UTR3 | ADAM17 | rs573141533 | PASS | vSR | 0 | 2342 | 6 | 0.002562 |
| **1** | 2 | 9488534 | A | G | UTR3 | ADAM17 | rs62120347 | PASS | vSR | 1 | 2342 | 53 | 0.02263 |
| **2** | 2 | 9488564 | A | G | UTR3 | ADAM17 | rs1052220677 | PASS | vSR | 0 | 2342 | 1 | 0.000427 |
| **3** | 2 | 9488580 | A | G | UTR3 | ADAM17 | rs192592002 | PASS | vSR | 0 | 2342 | 1 | 0.000427 |
| **4** | 2 | 9488645 | C | T | UTR3 | ADAM17 | rs899526914 | PASS | vSR | 0 | 2342 | 1 | 0.000427 |
| **2739** | 21 | 41508030 | C | A | UTR5 | TMPRSS2 | rs565468881 | PASS | vSR | 0 | 2342 | 2 | 0.000854 |
| **2740** | 21 | 41508115 | - | CCTCCG | UTR5 | TMPRSS2 | rs543548317 | PASS | vSR | 0 | 2340 | 19 | 0.00812 |
| **2741** | 21 | 41508116 | CCTCCGCCTCCG | - | UTR5 | TMPRSS2 | rs912333018 | PASS | vSR | 0 | 2304 | 1 | 0.000434 |
| **2742** | 21 | 41508131 | C | G | UTR5 | TMPRSS2 | rs561266954 | PASS | vSR | 0 | 2342 | 4 | 0.001708 |
| **2743** | 21 | 41508158 | C | A | UTR5 | TMPRSS2 | rs146132480 | PASS | vSR | 0 | 2342 | 25 | 0.010675 |

Figure 3.2: PyABraOM gene query batch search result for *ADAM17* and *TMPRSS2* on ABraOM database. The first and last five rows from the data table for *ADAM17* and *TMPRSS2*, respectively, is demonstrated.

#### 3.3 BIOVARS

BIOVARS was developed to aid users to access information from different human genetic variant databases in a fast and programmatic approach. The users can define which

[2] https://github.com/bioinfo-hcpa/pyABraOM

databases are to be considered when performing a variant search with the package. Although BIOVARS gets data for one or multiple queries across different data sources, some types of variant searches are not supported by every database and can throw an error if the user tries to perform such searches. One example is the search for variants given a specific transcript (gnomAD covers this functionality, but ABraOM can not handle such search operation). To demonstrate the BIOVARS functionalities, we will show how to get information from gnomAD and ABraOM databases, as well as possible search configurations for retrieving variant data for *ACE2*, *ADAM17* and *TMPRSS2* genes.

The BIOVARS gene search result comprehends the data merged from gnomAD and ABraOM databases. The table provides variant identifier, variants' annotation, genome localization, allele frequencies for each human population. To note, the Variant ID is a string composed of the chromosome, the genomic region and the reference and alternative allele to merge data. It is used as index to merge data from both human variant databases. In the code example below, the *biovars_df_1* object is a table with 18 columns of information for 5837 variants according to the hg38 genome version using the BIOVARS batch search functionality. Some details of this table are shown in Figure 3.3.

```
# Import modules for performing the search operations
from biovars import Sources, Search

# Prepare inputs of different search types
gene_list =["ACE2","ADAM17","TMPRSS2"]
region_list = [("X-15562032-15564176"),("2-9490303-9493719"),
               ("21-41480500-41480570")]
t_list = ["ENST00000427411","ENST00000310823","ENST00000332149"]

# Use both databases in the search
src = Sources(ref_genome_version="hg38", gnomad=True, abraom=True)

# Initialize the Search object
sch = Search(src)

# Perform the genes, regions and transcripts searches
biovars_df_1 = sch.gene_search(gene_list)
biovars_df_2 = sch.region_search(region_list)

biovars_df_3_1 = sch.transcript_search(t_list)
# An error raises: Transcript search not supported by ABraOM database!

# Disable ABraOM from the data sources to properly execute the search
src = Sources(ref_genome_version="hg38", gnomad=True, abraom=False)
sch = Search(src)
biovars_df_3_2 = sch.transcript_search(t_list)
```

| Variant ID | rsID | Gene | Annotation | Chromosome | Location | Reference | Alternative | African | Amish | Latino | Ashkenazi Jewish | East Asian | European (Finnish) | European (non-Finnish) | Other | South Asian | Middle Eastern | Brazilian ABraOM |
|---|---|---|---|---|---|---|---|---|---|---|---|---|---|---|---|---|---|---|
| **2-9488499-GTATT-G** | rs573141533 | ADAM17 | 3_prime_UTR_variant | 2 | 9488499 | GTATT | G | 0.000000 | 0.0 | 0.000000 | 0.0 | 0.00000 | 0.0 | 0.000000 | 0.0 | 0.000000 | 0.0 | 0.002562 |
| **2-9488534-A-G** | rs62120347 | ADAM17 | 3_prime_UTR_variant | 2 | 9488534 | A | G | 0.000000 | 0.0 | 0.000000 | 0.0 | 0.00000 | 0.0 | 0.000000 | 0.0 | 0.000000 | 0.0 | 0.022630 |
| **2-9488564-A-G** | rs1052220677 | ADAM17 | 3_prime_UTR_variant | 2 | 9488564 | A | G | 0.000000 | 0.0 | 0.000000 | 0.0 | 0.00000 | 0.0 | 0.000000 | 0.0 | 0.000000 | 0.0 | 0.000427 |
| **2-9488580-A-G** | rs192592002 | ADAM17 | 3_prime_UTR_variant | 2 | 9488580 | A | G | 0.000000 | 0.0 | 0.000000 | 0.0 | 0.00000 | 0.0 | 0.000000 | 0.0 | 0.000000 | 0.0 | 0.000427 |
| **2-9488645-C-T** | rs899526914 | ADAM17 | 3_prime_UTR_variant | 2 | 9488645 | C | T | 0.000000 | 0.0 | 0.000000 | 0.0 | 0.00000 | 0.0 | 0.000000 | 0.0 | 0.000000 | 0.0 | 0.000427 |
| **X-15600935-A-G** | rs774977446 | ACE2 | 5_prime_UTR_variant | X | 15600935 | A | G | 0.000000 | 0.0 | 0.000000 | 0.0 | 0.00000 | 0.0 | 0.000019 | 0.0 | 0.000000 | 0.0 | 0.000000 |
| **X-15600945-T-C** | rs370596467 | ACE2 | 5_prime_UTR_variant | X | 15600945 | T | C | 0.000000 | 0.0 | 0.000095 | 0.0 | 0.00168 | 0.0 | 0.000000 | 0.0 | 0.000375 | 0.0 | 0.000000 |
| **X-15600948-C-T** | rs184503057 | ACE2 | 5_prime_UTR_variant | X | 15600948 | C | T | 0.000033 | 0.0 | 0.000000 | 0.0 | 0.00000 | 0.0 | 0.000000 | 0.0 | 0.000000 | 0.0 | 0.000000 |
| **X-15600977-A-G** | rs1012928517 | ACE2 | 5_prime_UTR_variant | X | 15600977 | A | G | 0.000000 | 0.0 | 0.000000 | 0.0 | 0.00000 | 0.0 | 0.000000 | 0.0 | 0.000000 | 0.0 | 0.000000 |
| **X-15600982-C-G** | rs1452729298 | ACE2 | 5_prime_UTR_variant | X | 15600982 | C | G | 0.000000 | 0.0 | 0.000000 | 0.0 | 0.00000 | 0.0 | 0.000038 | 0.0 | 0.000000 | 0.0 | 0.000000 |
| **21-41464550-G-A** | rs76000363 | TMPRSS2 | downstream | 21 | 41464550 | G | A | 0.000000 | 0.0 | 0.000000 | 0.0 | 0.00000 | 0.0 | 0.000000 | 0.0 | 0.000000 | 0.0 | 0.084543 |
| **21-41464558-GA-G** | rs143680939 | TMPRSS2 | 3_prime_UTR_variant | 21 | 41464558 | GA | G | 0.000000 | 0.0 | 0.000000 | 0.0 | 0.00000 | 0.0 | 0.000000 | 0.0 | 0.000000 | 0.0 | 0.084188 |
| **21-41464558-G-GA** | | TMPRSS2 | 3_prime_UTR_variant | 21 | 41464558 | G | GA | 0.000000 | 0.0 | 0.000000 | 0.0 | 0.00000 | 0.0 | 0.000000 | 0.0 | 0.000000 | 0.0 | 0.000000 |
| **21-41464569-T-C** | rs456142 | TMPRSS2 | 3_prime_UTR_variant | 21 | 41464569 | T | C | 0.000000 | 0.0 | 0.000000 | 0.0 | 0.00000 | 0.0 | 0.000000 | 0.0 | 0.000000 | 0.0 | 0.767293 |
| **21-41464648-T-C** | rs948202058 | TMPRSS2 | 3_prime_UTR_variant | 21 | 41464648 | T | C | 0.000000 | 0.0 | 0.000000 | 0.0 | 0.00000 | 0.0 | 0.000000 | 0.0 | 0.000000 | 0.0 | 0.000427 |

Figure 3.3: BIOVARS gene query batch search result for *ACE2*, *ADAM17* and *TMPRSS2* on gnomAD and ABraOM database.The five first rows of *ACE2* and *ADAM17* and five last rows from the data table for *TMPRSS2* are demonstrated. To note, the Variant ID column is an indexer to integrate data from both databases created by BIOVARS package.

CHAPTER 4

### Graphical Visualization

Graphical visualization is a powerful data-driven analytics method widely used to increase the comprehension over the investigated data. It is of particular interest when dealing with large data sets and their imminent complexity to the human eye. Thus, additionally to APIs, we provide plotting methods coded in R (interfaced by the *rpy2* Python package) for summarizing the variants search results from gnomAD and ABraOM databases. Both static and interactive visualizations of the results are provided by BIOVARS and these methods can also be applied to Pynoma and PyABraOM resulting dataframes, as long as they utilize a helper method to convert their specific dataframes to the standard BIOVARS's expected format. Through the implemented visualizations, the user can visualize allele frequency and annotation of a given region and see the position where variants fall inside canonical and non canonical transcripts by using annotations from the Ensembl database for both latest human versions (hg37 and hg38). Besides, the total, shared and private variant count among human populations is provided on statistical plots. The interactive visualization of the results is given in an HTML file named Variant Report, which enables the appraisal of the data quantitatively and qualitatively. In the remaining of this Chapter, we demonstrate how to use the visualization method for the example of *ACE2* from section 2.2 and the genomic region X:15562032-15564176.

#### 4.1 Load libraries for graphical visualization

```
# Load all libraries for plot visualizations
from biovars import Plotter
```

#### 4.2 Total, private and shared variants charts

The possible types of visualization can be used in terms of private, shared and total population variants where the user can plot the data on a map Figure 4.1 or only plot the bar plots in a grid Figure 4.2 for a considered frequency threshold. If no frequency is set, the plot will contain variants with frequency equal or greater than 1% in populations. Furthermore, the *saving_path* parameter must be indicated with the path and the name of the file followed by the user's desired extension - which has to be either *.pdf* or *.png*.

```
# Input
ace2_data = biovars_df_1[biovars_df_1["Gene"] =="ACE2"]

# Initialized  an object
plt = Plotter(dataframe=ace2_data, genome_version="hg38")

# Plot world map by using the 1% threshold frequency
plt.plot_world(saving_path="/home/user/path/world_map.png", frequency
    =0.01)
```

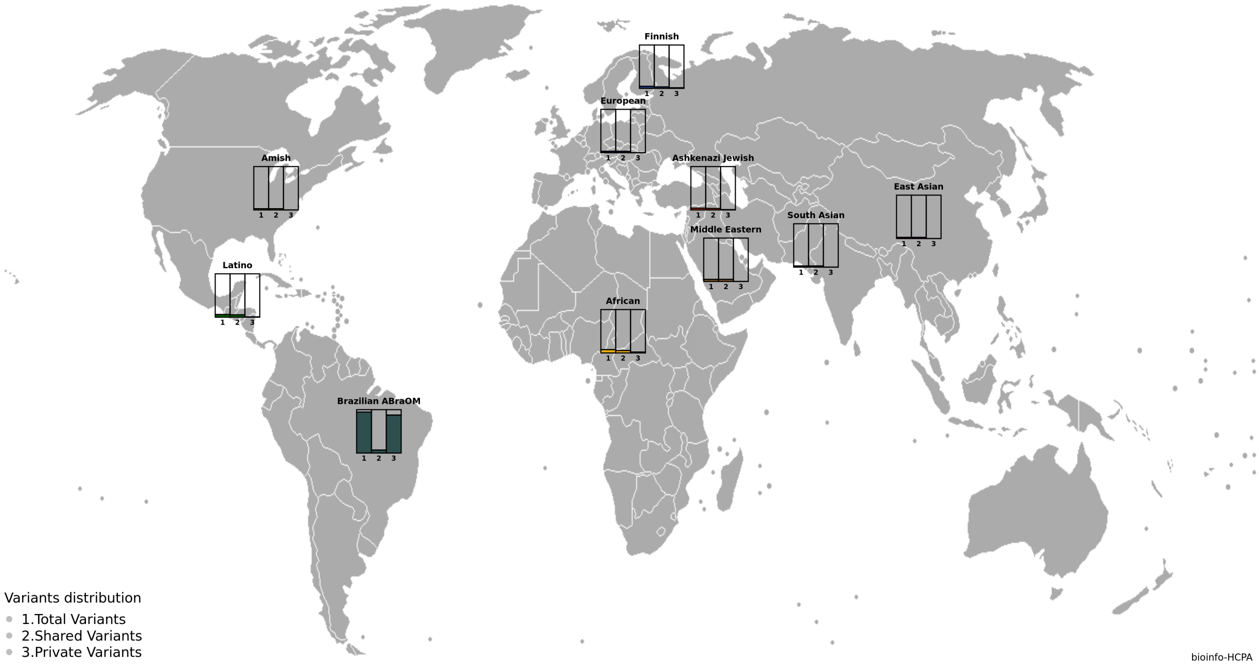

Figure 4.1: World map visualization with total, shared and private variants bar plot with frequency equal or grater than 1% among human populations for *ACE2* gene.

```
# Plot only the grid with the bar plots by using the 1% threshold
    frequency
plt.plot_variants_grid(saving_path="/home/user/path/grid_bar_plot.png",
    frequency=0.01)
```

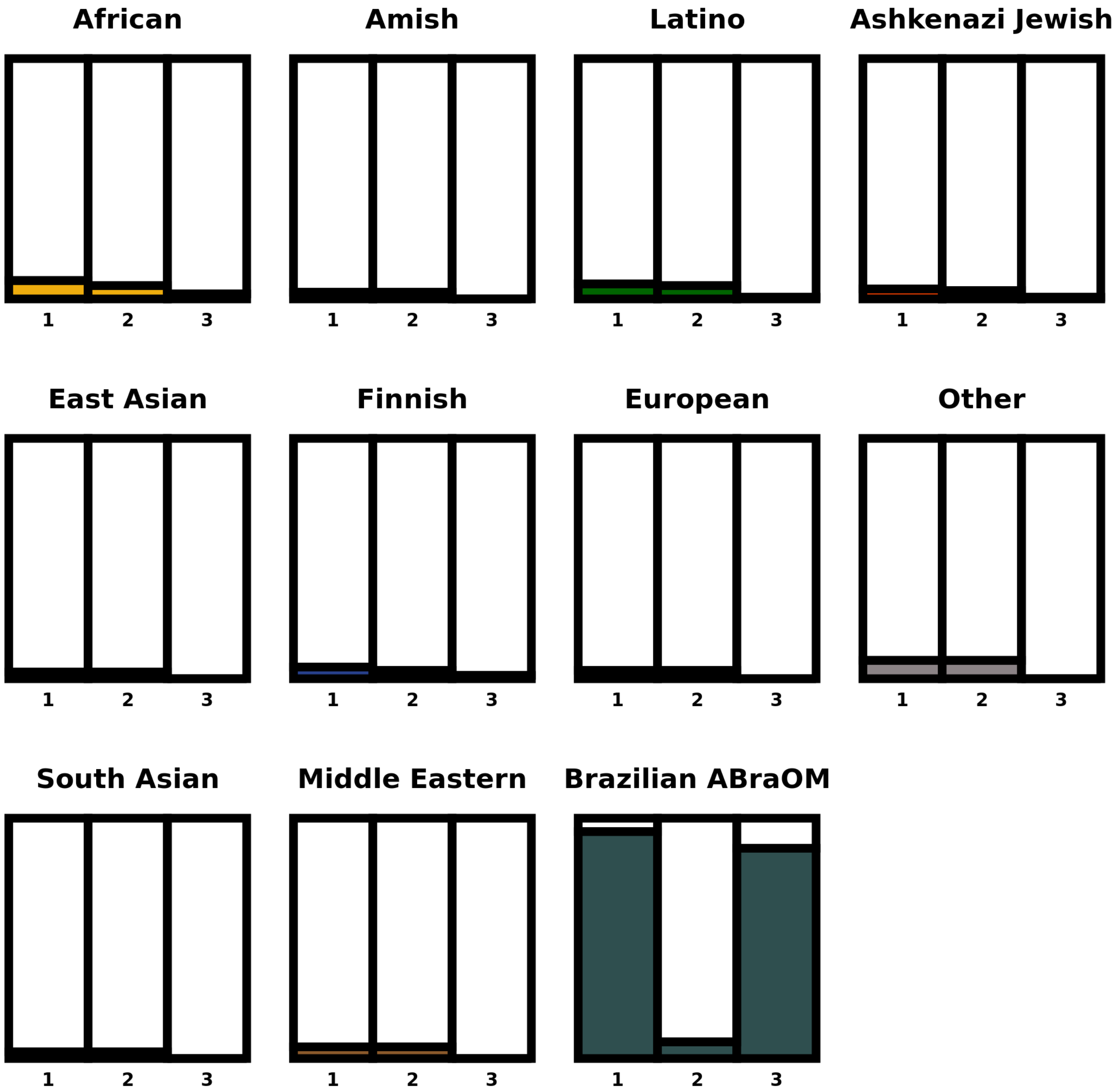

Figure 4.2: Grid bar plot of total, shared and private variants with frequency equal or grater than 1% among human populations for *ACE2* gene.

#### 4.3 Variant Frequency and Annotation

Plots the genomic region within the specified start and end range (max. of 54bp) with the transcripts and location, frequency (Figure 4.3 and Figure 4.4 ) and annotation (Figure 4.5 and Figure 4.6) for each type of variant found in the data frame along the specified region. This region must be contained inside the Plotter data frame. To note, the localization of variants into the transcripts for variant frequency and annotation can be represented by considering a specific region or throughout the transcripts length for Canonical and Non Canonical transcripts.

```
#The variant allele frequencies are indicated in the plots.
plt.plot_genomic_region(saving_path="/home/user/path/allele_frequency1.
    pdf",starting_region=15562032, ending_region=15564176, mut=False,
    transcript_region = True)
```

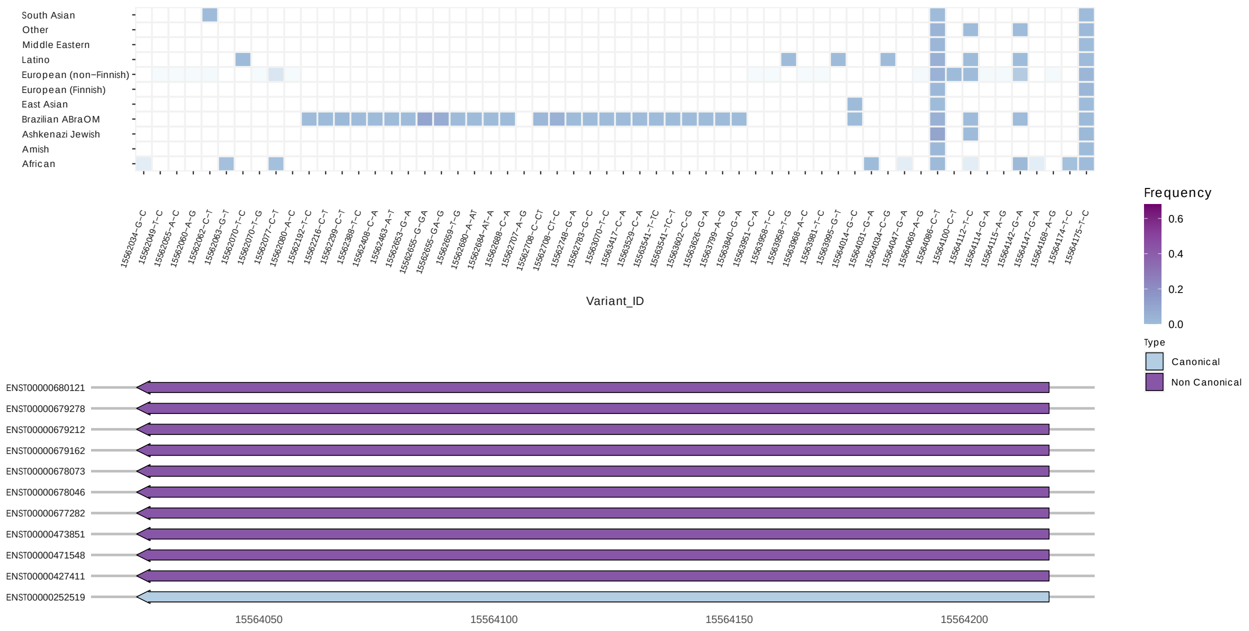

Figure 4.3: Chart of *ACE2* frequency variants in the genome region X:15562032-15564176. (A) Grid of variant frequencies among human populations. (B) The specific region of canonical and Non Canonical transcripts region where the variants are located.

```
#The variant allele frequencies are indicated in the plots and signed to
    a region in transcripts
plt.plot_genomic_region(saving_path= "/home/user/path/allele_frequency2.
    pdf", starting_region= 15562032, ending_region= 15564176, mut= False,
     transcript_region= False)
```

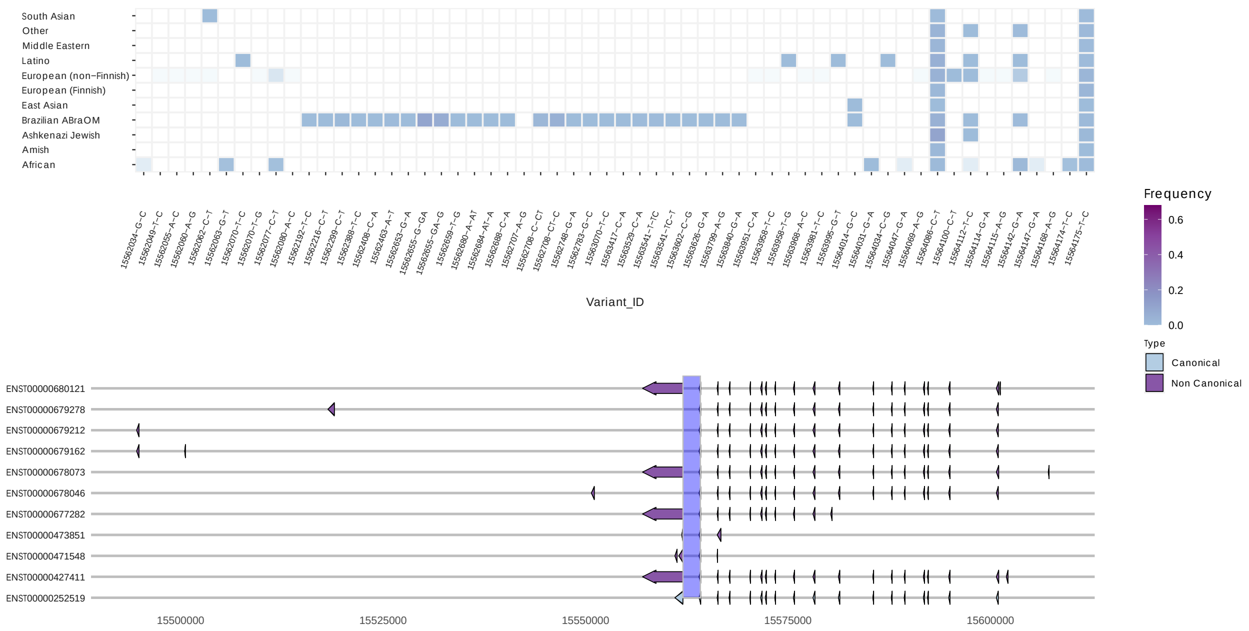

Figure 4.4: Chart of *ACE2* frequency variants in the genome region X:15562032-15564176. (A) Grid of variant frequencies among human populations. (B) Canonical and Non Canonical transcript gene information. Vertical line indicates the region of variant are located.

```
# Set transcript_region to True for showing where the variant falls inside a specific region inside the transcripts.
plt.plot_genomic_region(saving_path="/home/user/path/variant_annotation1.pdf",starting_region = 15562032, ending_region = 15564176, mut = True, transcript_region = True)
```

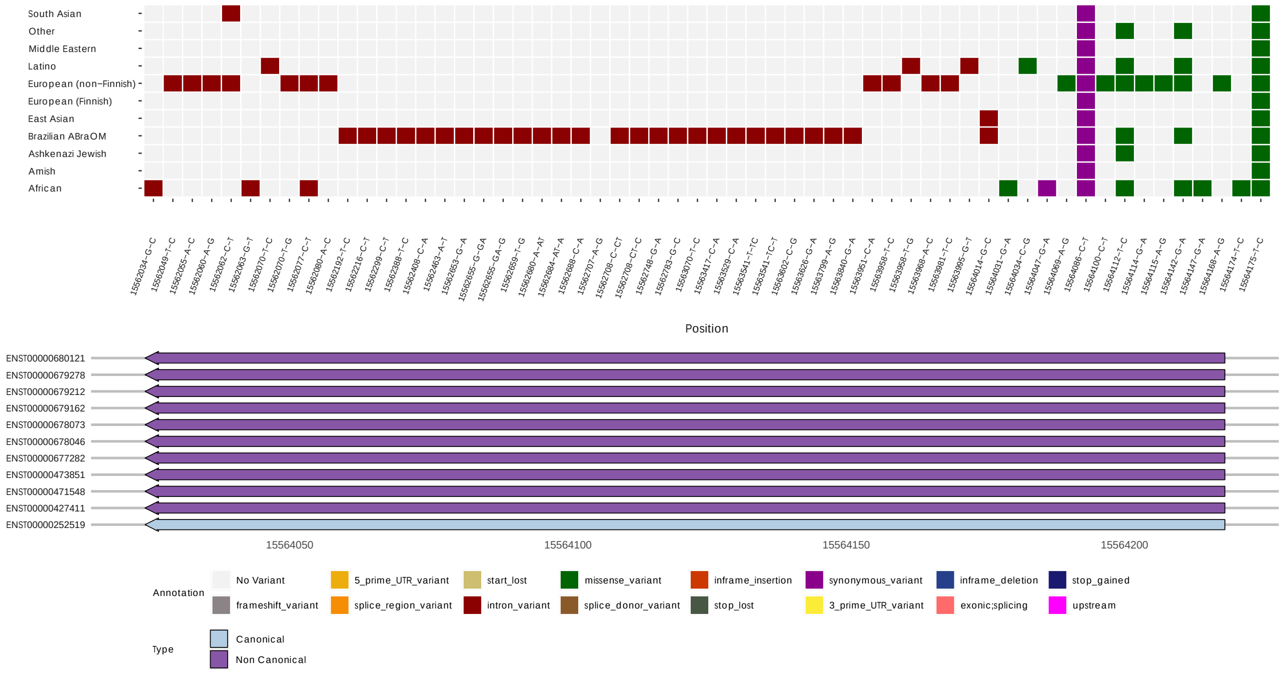

Figure 4.5: Chart of variant annotations across genome region X:15562032-15564176 of *ACE2* gene. (A) Grid of variant annotation among human populations.(B) The specific region of canonical and Non Canonical transcripts region where the variants are localized. The legend below represent the variant color annotation and transcript type on the charts.

```
# Set transcript_region to False for showing where the variant falls inside all transcript length.
plt.plot_genomic_region(saving_path="/home/user/path/variant_annotation2.pdf", starting_region=15562032, ending_region=15564176, mut=True, transcript_region=False)
```

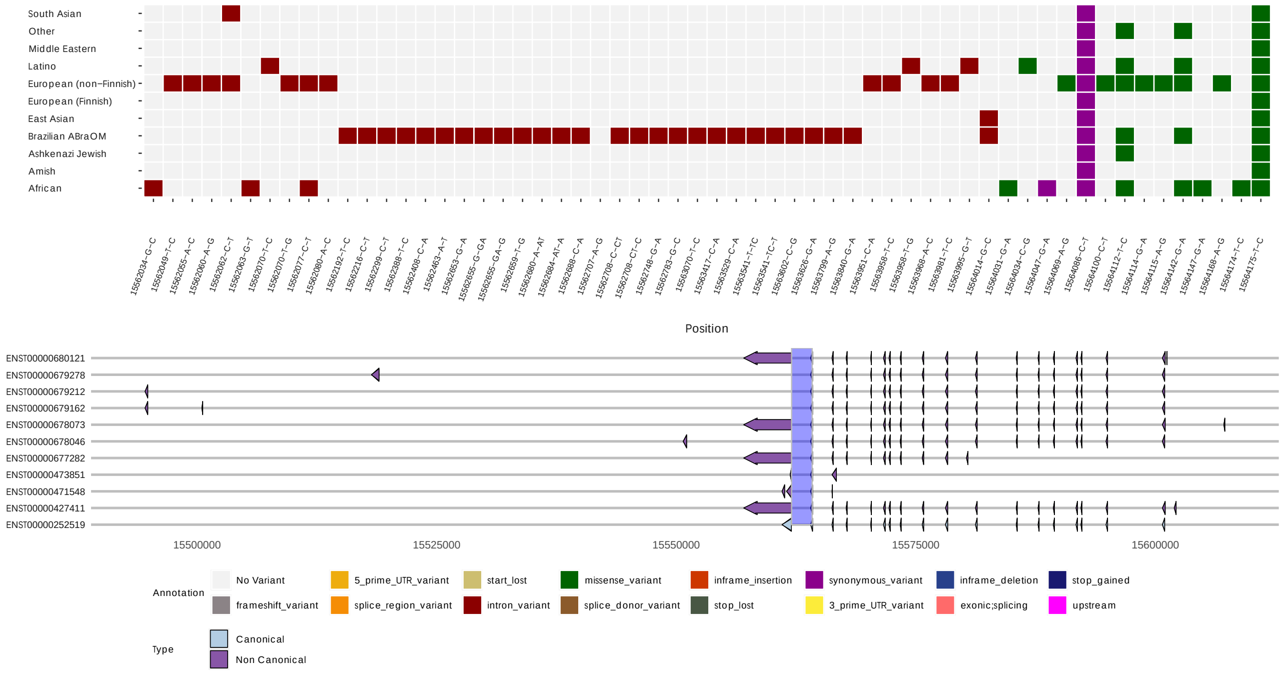

Figure 4.6: Chart of *ACE2* frequency variants in the genome region X:15562032-15564176. (A) Grid of variant frequencies among human populations. (B) Canonical and Non Canonical transcript gene information. Vertical line indicates the region of variant are localized inside each transcript. The legend below represent the variant color annotation and transcript type on the charts.

Some adjustments could be made to use the same function to plot the frequency or variant annotation by changing *mut* parameter, which is used to inform whether it should be indicated in the plots. Besides, the *transcript_region* parameter can be set to visualize where the variants are located by considering a specific region or throughout the transcripts length. Furthermore, the *saving_path* parameter must be indicated with the path and the name of the file followed by the user's desired extension - which has to be either *.pdf* or *.png*

#### 4.4 Interactive visualization

The code below generates a variant report in HTML file format inside the chosen directory. It contains all plots mentioned above (except for the variant grid) along with plotted data frame for interactive view to perform data quantitative and qualitative analysis.

The interactive table in Figure 4.5 shows the input data provided in the object initializer (see Section 4.2). This visualization provides filters that optimize data visualization in the data table. The following part of Variant report shows a bar plot of variants present in human populations, as shown in Figure 4.8. The expansion of pop-ups show the total, private and shared variants for each population based on the variant frequency threshold. The interactive variant frequency in Figure 4.9 and annotation in Figure 4.10 charts shows the information for the genomic region chrX:15573310-15573460 from *ACE2* gene. The information of frequency and annotation is displayed by color and value inside the pop-up. Gray color indicates variants that are not find in the specific population. Canonical and Non Canonical transcript details are displayed for those variants located inside genes, as seen in Figure 4.11.

```
plt.plot_summary(saving_directory = "/home/user/path/",gene = "ACE2",
    starting_region =15573310,ending_region = 15573460,  frequency =
    0.01)
```

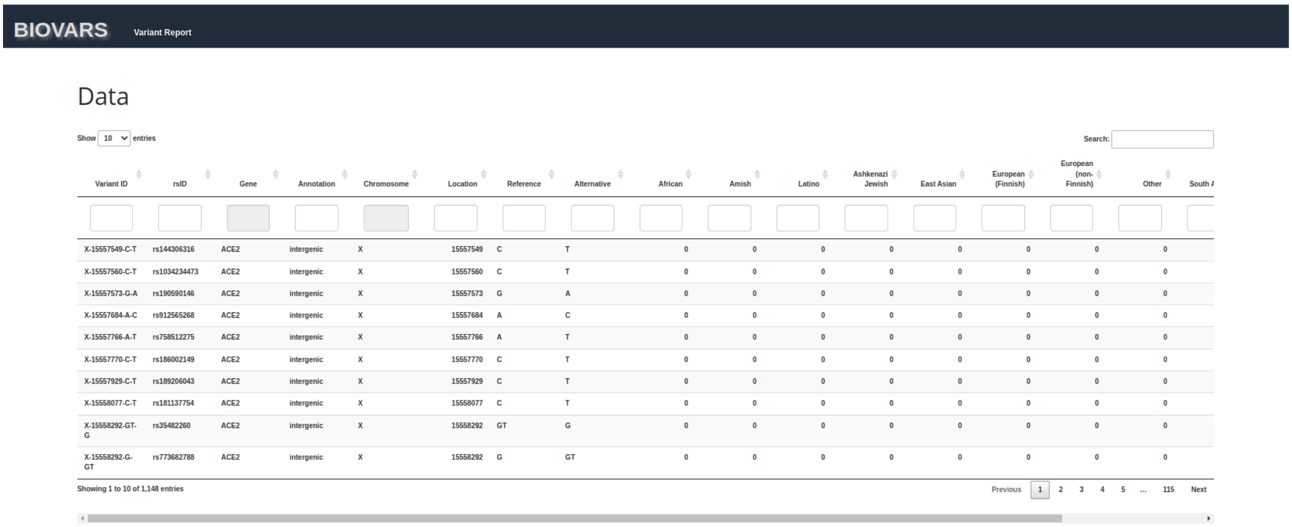

| Variant ID | rsID | Gene | Annotation | Chromosome | Location | Reference | Alternative | African | Amish | Latino | Ashkenazi Jewish | East Asian | European (Finnish) | European (non-Finnish) | Other | South A |
|---|---|---|---|---|---|---|---|---|---|---|---|---|---|---|---|---|
| X-15557549-C-T | rs144306316 | ACE2 | intergenic | X | 15557549 | C | T | 0 | 0 | 0 | 0 | 0 | 0 | 0 | 0 | |
| X-15557560-C-T | rs1034234473 | ACE2 | intergenic | X | 15557560 | C | T | 0 | 0 | 0 | 0 | 0 | 0 | 0 | 0 | |
| X-15557573-G-A | rs190590146 | ACE2 | intergenic | X | 15557573 | G | A | 0 | 0 | 0 | 0 | 0 | 0 | 0 | 0 | |
| X-15557684-A-C | rs912565268 | ACE2 | intergenic | X | 15557684 | A | C | 0 | 0 | 0 | 0 | 0 | 0 | 0 | 0 | |
| X-15557766-A-T | rs758512275 | ACE2 | intergenic | X | 15557766 | A | T | 0 | 0 | 0 | 0 | 0 | 0 | 0 | 0 | |
| X-15557770-C-T | rs186002149 | ACE2 | intergenic | X | 15557770 | C | T | 0 | 0 | 0 | 0 | 0 | 0 | 0 | 0 | |
| X-15557929-C-T | rs189206043 | ACE2 | intergenic | X | 15557929 | C | T | 0 | 0 | 0 | 0 | 0 | 0 | 0 | 0 | |
| X-15558077-C-T | rs181137754 | ACE2 | intergenic | X | 15558077 | C | T | 0 | 0 | 0 | 0 | 0 | 0 | 0 | 0 | |
| X-15558292-GT-G | rs35482260 | ACE2 | intergenic | X | 15558292 | GT | G | 0 | 0 | 0 | 0 | 0 | 0 | 0 | 0 | |
| X-15558292-G-GT | rs773682788 | ACE2 | intergenic | X | 15558292 | G | GT | 0 | 0 | 0 | 0 | 0 | 0 | 0 | 0 | |

Figure 4.7: BIOVARS data table from Variant Report HTML file for *ACE2* gene. The human genetic variants information and the fields to filter to easy data manipulation are displayed.

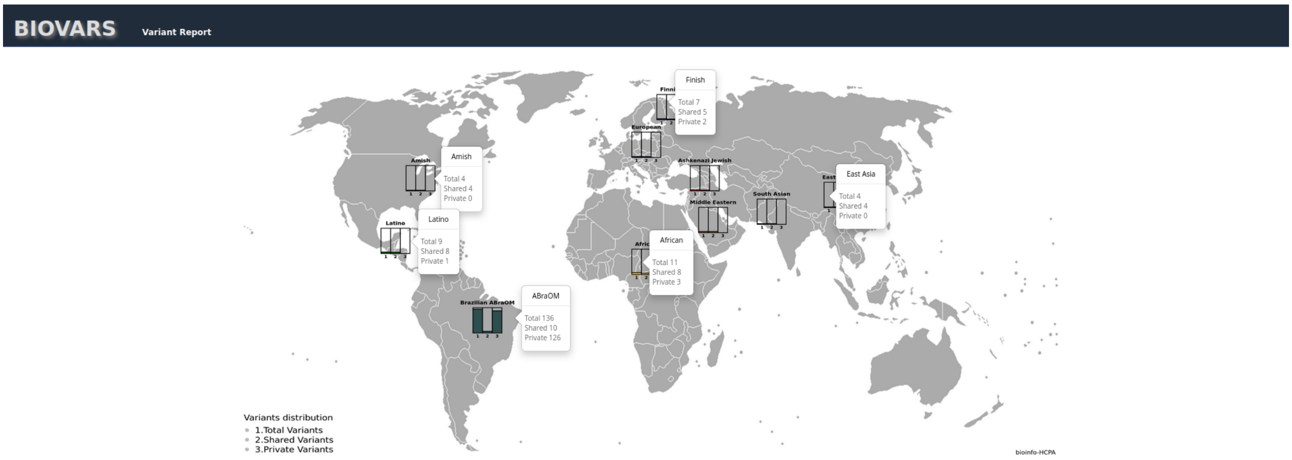

Figure 4.8: Number of the total, shared, and private variants among different human populations from Variant Report HTML file for *ACE2* gene. The pop-up expansion shows the number of variants for each population in their respectively category.

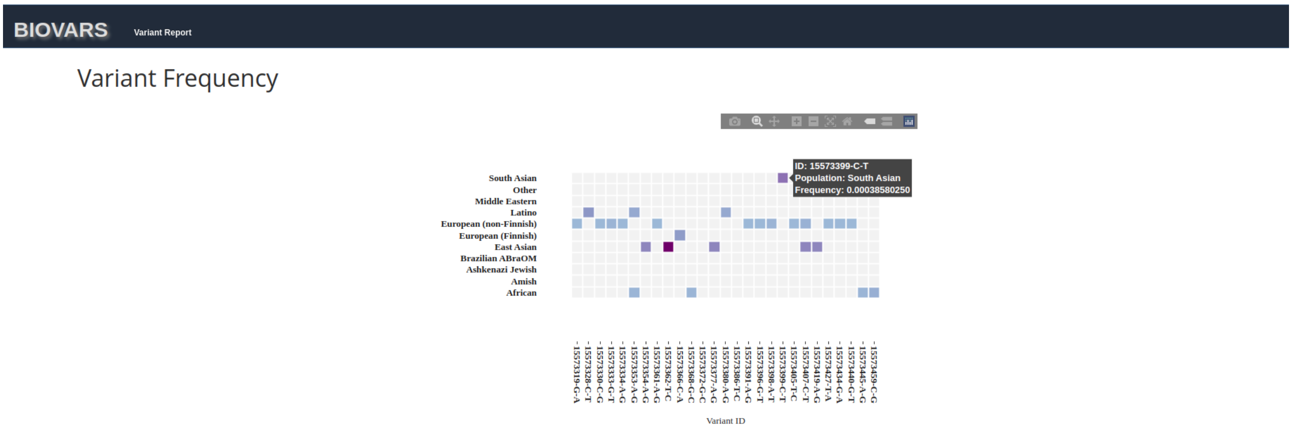

Figure 4.9: Variant frequency information from Variant Report HTML file for *ACE2* gene. The allele frequency for variants inside the chrX:15573310:15573460 among human population are shown.

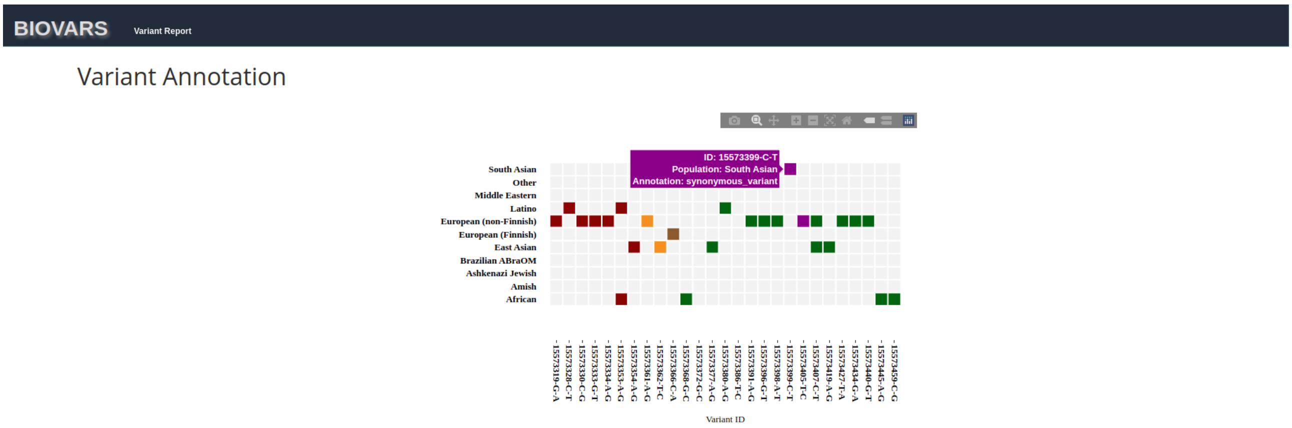

Figure 4.10: Variant annotation from Variant Report HTML file for *ACE2* gene.The variants for genomic region chrX:15573310:15573460 in human populations are shown.

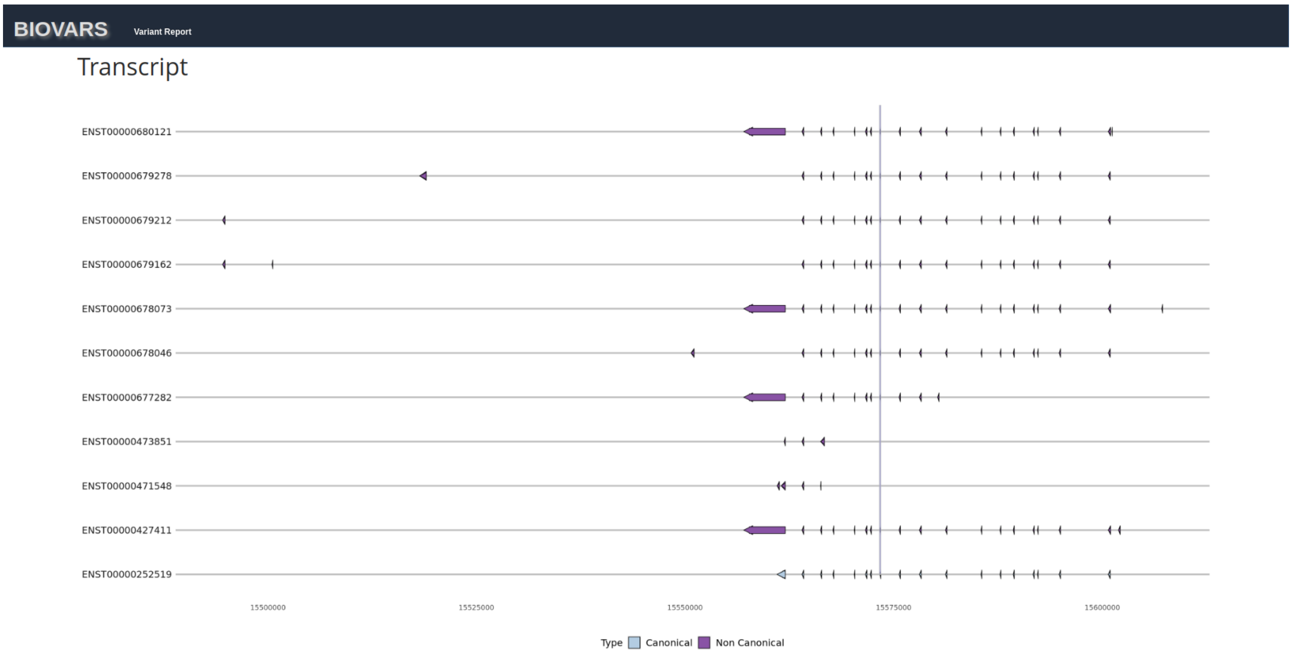

Figure 4.11: Transcript information from Variant Report HTML file for *ACE2* gene. Canonical and Non Canonical transcript gene information. Vertical line indicates the position where the variant is located in each transcript.

### References

I. Glowacka, S. Bertram, M. A. Müller, P. Allen, E. Soilleux, S. Pfefferle, I. Steffen, T. S. Tsegaye, Y. He, K. Gnirss, et al. Evidence that tmprss2 activates the severe acute respiratory syndrome coronavirus spike protein for membrane fusion and reduces viral control by the humoral immune response. *Journal of virology*, 85(9):4122–4134, 2011.

K. J. Karczewski, L. C. Francioli, G. Tiao, B. B. Cummings, J. Alföldi, Q. Wang, R. L. Collins, K. M. Laricchia, A. Ganna, D. P. Birnbaum, et al. The mutational constraint spectrum quantified from variation in 141,456 humans. *Nature*, 581(7809):434–443, 2020.

D. W. Lambert, M. Yarski, F. J. Warner, P. Thornhill, E. T. Parkin, A. I. Smith, N. M. Hooper, and A. J. Turner. Tumor necrosis factor-$\alpha$ convertase (adam17) mediates regulated ectodomain shedding of the severe-acute respiratory syndrome-coronavirus (sars-cov) receptor, angiotensin-converting enzyme-2 (ace2). *Journal of Biological Chemistry*, 280(34): 30113–30119, 2005.

M. S. Naslavsky, M. O. Scliar, G. L. Yamamoto, J. Y. T. Wang, S. Zverinova, T. Karp, K. Nunes, J. R. M. Ceroni, D. L. de Carvalho, C. E. da Silva Simões, et al. Whole-genome sequencing of 1,171 elderly admixed individuals from são paulo, brazil. *Nature communications*, 13(1): 1–11, 2022.
